# Dynamic coupling of residues within proteins as a mechanistic foundation of many enigmatic pathogenic missense variants

**DOI:** 10.1101/2021.09.24.461738

**Authors:** Nicholas J. Ose, Brandon M. Butler, Avishek Kumar, Maxwell Sanderford, Sudhir Kumar, S. Banu Ozkan

**Affiliations:** Department of Physics and Center for Biological Physics, Arizona State University, Tempe, Arizona, 85281; Institute for Genomics and Evolutionary Medicine, Temple University, Philadelphia, PA 19122; Department of Biology, Temple University, Philadelphia, PA 19122; Center for Genomic Medicine and Research, King Abdulaziz University, Jeddah, Saudi Arabia

## Abstract

Many pathogenic missense mutations are found in protein positions that are neither well-conserved nor belong to any known functional domains. Consequently, we lack any mechanistic underpinning of dysfunction caused by such mutations. We explored the disruption of allosteric dynamic coupling between these positions and the known functional sites as a possible mechanism for such mutations. In this study, we present an analysis of 144 human enzymes containing 591 pathogenic missense variants, in which allosteric dynamic coupling of mutated positions with known active sites provides insights into a primary biophysical mechanism and evidence of their functional importance. We illustrate this mechanism in a case study of *β-Glucocerebrosidase* (GCase), which contains 94 Gaucher disease-associated missense variants located some distance away from the active site. An analysis of the conformational dynamics of GCase suggests that mutations on these distal sites cause changes in the flexibility of active site residues despite their distance, indicating a dynamic communication network throughout the protein. The disruption of the long-distance dynamic coupling due to the presence of missense mutations may provide a plausible general mechanistic explanation for biological dysfunction and disease.

**Author Summary:** Genetic diseases occur when mutations to a particular gene cause a gain/loss in function of the related protein. Although several methods based on conservation and protein biochemistry exist to predict which genetic mutations may impact function, many disease causing changes remain unexplained by these metrics. In this study, we propose an explanation for these genetic changes may cause disease. In order to function, important regions of a protein must be able to exhibit collective motion. Through computer simulations, we observed that changing even a single amino acid within a protein can change the protein motion. Notably, disease causing genetic changes tend to alter the motion of regions which are critically important to protein function, even the mutations are far from these critical regions. In addition, we examined the degree that two amino acids within a protein may “couple” to one another, meaning the degree to which motion in one amino acid will affect the other. We found that amino acids which are highly coupled to the active site of a protein are more likely to result in disease if mutated, thereby offering a new tool for predicting genetic disease which incorporates internal protein dynamics.

## Introduction

Our understanding of disease-causing genetic mutations continues to evolve. From a biophysics perspective, it has been shown that disease-associated variants could alter the stability of a protein [1–3]. Conversely, a high-throughput functional assay revealed that only one-third of over 2,000 mutations led to a decrease in protein stability [4]. Rather than affecting stability, a large fraction of disease-associated variants seems to impair specific protein-ligand function or enzymatic activity [5–8]. Furthermore, studies combining evolutionary approaches with the biochemistry of protein design have revealed that disease-causing mutations at non-conserved sites can involve complex and poorly understood mechanisms [5,9–11]. Through sequencing efforts, a large catalog of missense variants in thousands of human proteins has been assembled, including those implicated in diseases [5,11–13]. However, many pathogenic missense variants occur at positions that are neither evolutionary well-conserved nor a part of any known functional domain (Fig 1b-c). Regardless of biochemical similarity, amino acid substitutions at non-conserved sites lead to a wide range of outcomes, increasing or decreasing functional activity at up to three orders of magnitude (i.e., the rheostatic pattern of change) [14]. These enigmatic mutations are frequently misdiagnosed because neither evolutionary nor static structural features are informative. In fact, many rare missense variants occur at fast-evolving positions that do not have functional annotations (Fig 1c), which adversely impact the prediction accuracy of commonly used methods because they run counter to expectations (Fig 1d).

**Fig 1.**
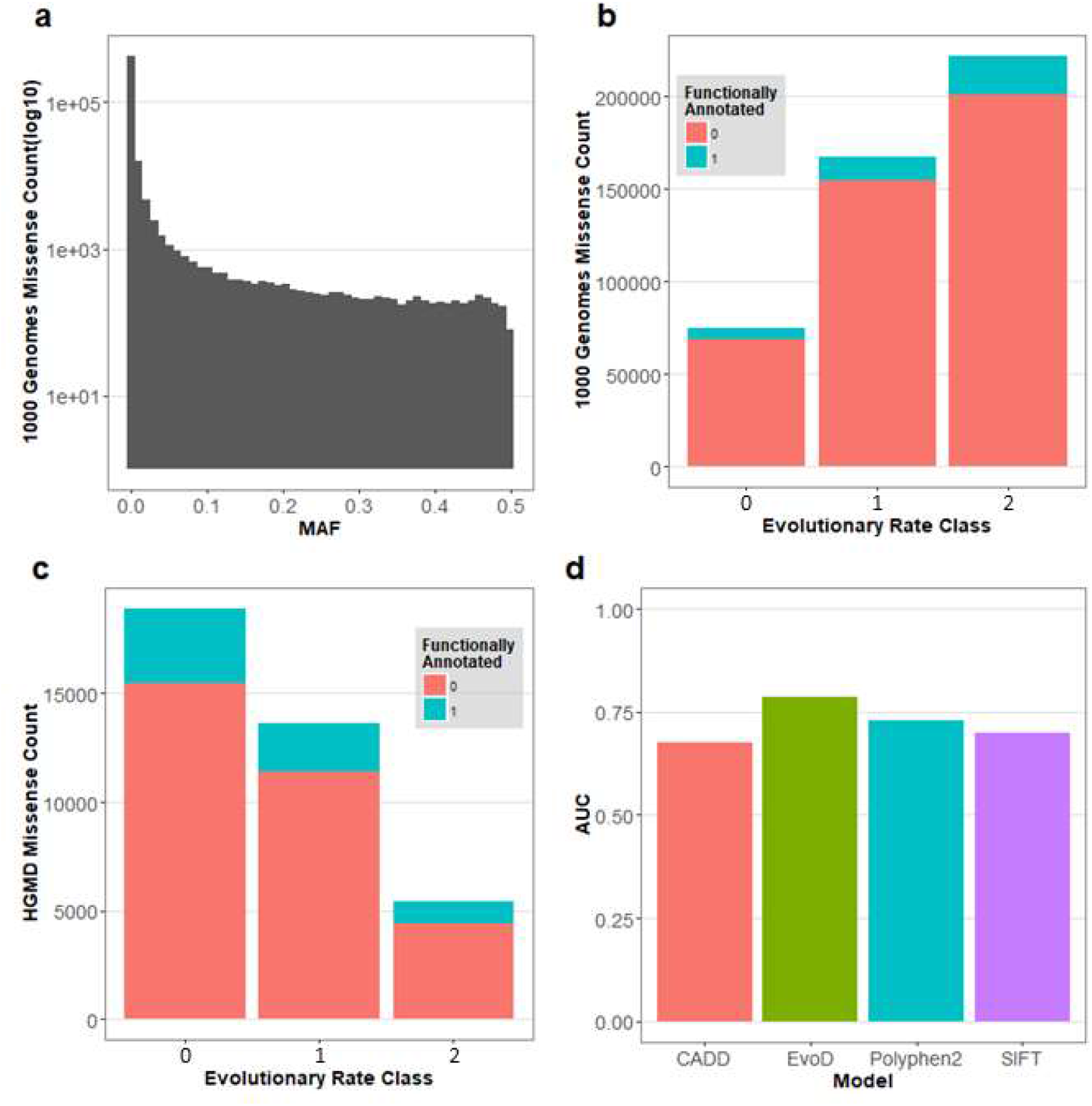
Frequency, evolutionary conservation, and rates of misdiagnosis of missense variants. (a) Histogram of minor allele frequencies (MAF) of missense variants in the 1000 Genomes data set. (b) Counts of these missense variants according to evolutionary conservation and the UNIPROT functional annotation of their domain of residence. Evolutionary rate classes are from Kumar et al. [15] with class 0 sites containing no substitutions, class 1 sites exhibiting 0-1 substitutions per billion years, and class 2 sites exhibiting greater than 1 substitution per billion years. (c) Histogram of evolutionary conservation of sites containing pathogenic missense variants with and without functional annotation in the Uniprot database. (d) Performance of four different missense diagnosis tools, quantified by their area under the receiver operation curve (AUC), which measures their ability to discriminate between putatively neutral (1000 Genomes missense variants with MAF>1%) and pathogenic missense variants found in fast evolving positions (evolutionary rate class of 2). Pathogenic variants with MAF > 0.01% were excluded.

Here we explore the mechanistic role of dynamic allosteric coupling of sites carrying disease mutations with the catalytic sites important for enzymatic activity. Our exploration is based on the premise that many mutations alter conformational dynamics of proteins, shifting the distribution of the ensemble and protein function, including the emergence of new functions [10,16–20], adaption to different environments [21], and dysfunction [12,22].

We used the *dynamic coupling index* (DCI) to identify sites strongly coupled to active sites critical for function [23,24]. We refer to them as dynamic allosteric residue coupling (DARC) sites. A mutation at a DARC site is likely to influence conformational dynamics and allosteric regulation, making individuals carrying mutants of these sites highly susceptible to disease phenotypes.

This mechanism is elucidated by examining GCase, a signature human enzyme wcontaining 497 amino acids and more than 200 known disease-associated missense variants. These missense mutations lead to Gaucher disease (GD) [25] characterized by a dangerous buildup of lipids in certain organs. Genetic changes in GCase can lead to other health conditions as well, including Parkinson’s disease [26–29] and Dementia with Lewy bodies [29,30]. We investigated the mechanistic impact of these mutations on conformational dynamics and allosteric regulation [31]. In the following, we show that GD mutations disrupt allosteric regulation due to changes in dynamic flexibility around the catalytic sites, altering enzymatic activity essential for homeostasis. The disease mutation sites can be thought of as key DARC sites. To further explore the role of conformational dynamics and allostery in missense variants at a broader scale, we conducted a proteome-wide analysis of allosteric coupling for a set of enzymes. These analyses suggest that pathogenic variants are most abundant at DARC sites. We also present an analysis of DCI asymmetry, which measures the degree of symmetry in the dynamic coupling between two sites, revealing that mutations are likely to result in a loss of function if they occur at distal sites controlled by the active site, resulting in pathogenesis.

## Results

### Disease-associated mutations modify dynamics throughout the protein

GCase is a member of the family of glycoside hydrolases that use glutamates for hydrolyzing glucocerebrosidase into glucose and ceramide. Many amino acid variants of this enzyme are reported to cause Parkinson’s disease [26–29], Dementia with lewy bodies [29,30], and GD [32]. Using the crystal structure of GCase (<u>1ogs.pdb</u>) [33] and 97 GD associated variants [34], we calculated the three-dimensional distance between the mutation site and the active site (residues 235 and 340). A vast majority (87.6%) of GD pathogenic variants occur further than 10 Å from the nearest active site residue, making direct interactions implausible. This suggests the existence of a network of indirect interactions through which a mutation at a distal site can induce dynamic changes at other regions of the protein and, by extension, impact protein function. The behavior of residues within this network can be examined by using a structural dynamics-oriented approach.

We illustrate the approach using structural dynamics by considering the example of a single mutant protein site, *N370S*, as this variant is present with a high frequency (∼70%) in the Ashkenazi Jewish population and has been extensively studied [26,31]. We first calculated the structural flexibility profiles of residues using a position-specific dynamic flexibility index, or DFI (see methods section). A comparison between the DFI profiles of the wild type variant of GCase with the *N370S* mutation is shown in Fig. 2a. The DFI profile provides an estimate for the role of each residue in mediating structure-encoded dynamics. As for GCase, the DFI profile indicates significant shifts in dynamics caused by the *N370S* mutation. Regions of the protein that should be rigid are now flexible and vice versa (Fig 2b). Hinges of the protein have moved or disappeared, and new hinges have appeared elsewhere. As seen in previous studies, these hinge shifts suggest a major change in dynamics and thus, protein function [18,23,24,35].

**Fig 2.**
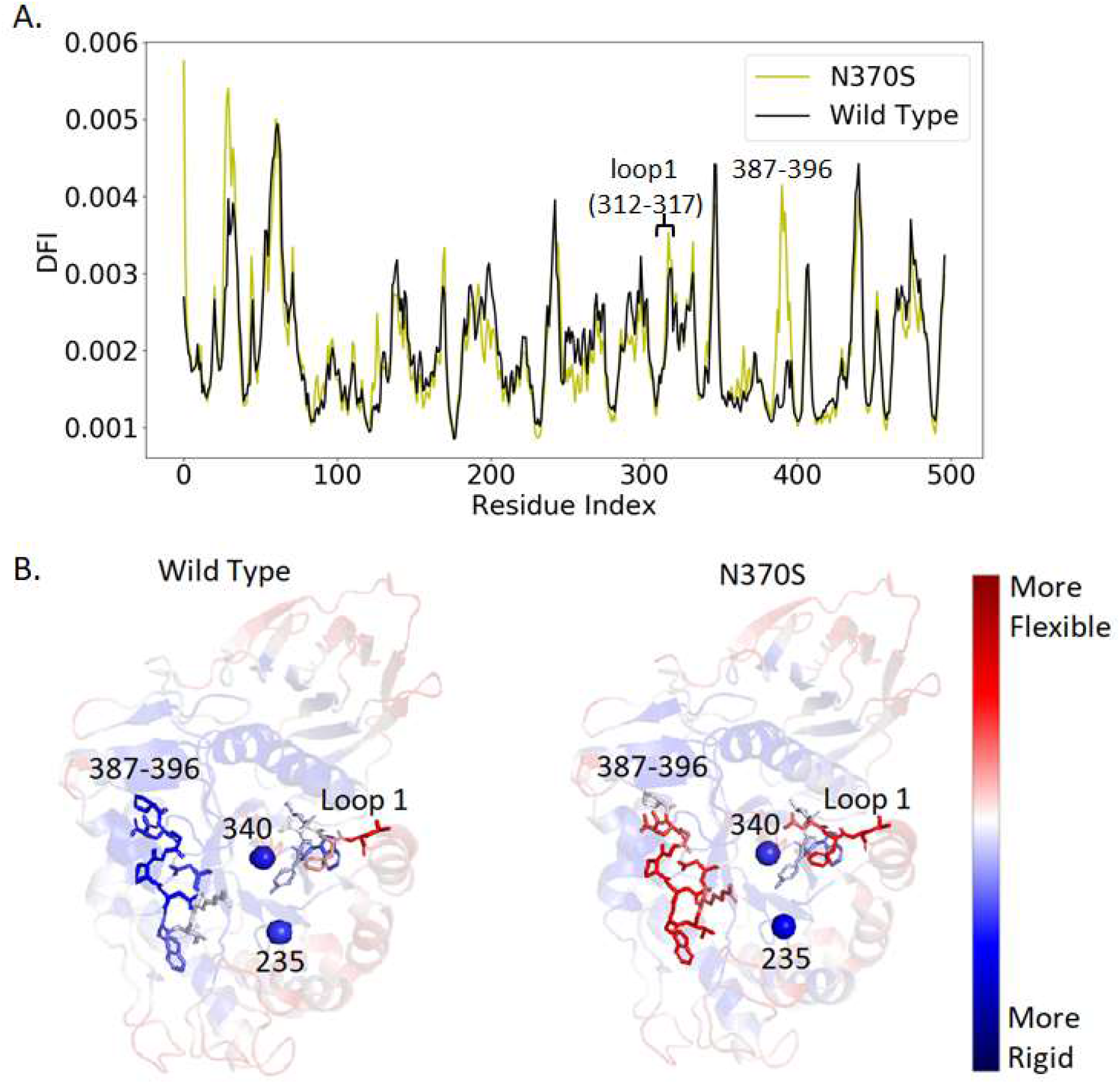
A comparison of N370S and wild type DFI profiles. (A) The %DFI profile of a mutant protein (*N370S*, yellow) contrasts with that of the wild type (black). The dissimilarities in the profiles of N370S and the wild type demonstrate how a single point disease mutation can induce changes in the flexibility profile of a different region of the protein. (B) Ribbon diagrams of both variants showing DFI as a color-coded spectrum from red-white-blue (red and blue indicates the highest and lowest flexibility, respectively). The regions with the most significant changes in dynamic flexibility are highlighted.

Among the five loops surrounding the active site, we observe that loop 1 (residues 312-317) exhibits an increase in DFI (Fig 2a), suggesting that increased flexibility of this loop could contribute to the decrease in enzymatic activity by hindering the accessibility of the ligand to the active site as observed in previous work [36]. This variant displays a small change in flexibility near loop 1. Changes in DFI within loop 1 for other studied mutations are shown in S1 Fig. Additionally, *N370S* shows a very large shift in flexibility between residues 387 to 396, which overlaps with loop 3 (residues 394-399); within the overlap is the *R395* residue, which orients differently in the active and inactive states of the enzyme [22] (Fig 2a).

### Mutations at distal sites dynamically-coupled to the active site alter long-range communication

Although the only sequence difference between the wild type and mutant GCase is a single residue, we observed changes in DFI across the protein. This suggests that changes in long-range dynamic coupling may be responsible for the altered flexibility profiles. The dynamic coupling index (DCI) captures the strength of a displacement response for position *i* upon perturbation of position *j*, relative to the average fluctuation response of position *i* to all other positions in the protein. In this way, DCI can reveal the degree of dynamic coupling between *i* and *j*.

Here, we present DCI as a percentile rank of the DCI range observed with values ranging from 0 to 1 (%DCI). Importantly, DFI and DCI are distinct in that DFI measures the flexibility of a position. In contrast, DCI measures the coupling of a position. Furthermore, DCI estimates are obtained in the context of the functional position selected for analysis. Every amino acid position in any given protein has a unique network of direct, local interactions that give rise to a unique network of highly coupled pair positions. Across the protein structure, this gives rise to an inhomogeneous 3D interaction network. Using DCI to observe this network is particularly useful when considering active sites because it is known that even far away positions may disrupt function through the mechanism of allostery [35]. Residues that are distant from the active site (>10 Å) yet are highly coupled to them (%DCI > 60) generally implies a greater than average response fluctuation when active site residues are perturbed) can play an important role in protein function.

In the example of GCase, around half (52.6%) of the studied pathogenic variants, including N370S, occur at DARC sites. In fact, according to our list of disease mutation sites [37], approximately 28% of DARC sites are associated with GD, compared to ∼15% of non-DARC sites throughout the entire protein. This suggests that the substitutions at DARC sites are more likely to lead to genetic disease. Moreover, a comparison of DCI values of sites with disease-associated missense variants with all other protein sites supports the same observation: mutations at DARC sites, distal sites that exhibit high coupling (i.e., high DCI), are predisposed to impact function [23,24].

When comparing DCI profiles of the active site for wild type and disease proteins, fluctuations in DCI occur at certain sites, while mostly decreasing around sites containing disease mutations (Fig 3c). These changes in DCI imply that the long-distance communication pathways cannot follow typical channels to the active site. This communication breakdown is presumed to be a consequence of altered dynamics. Losing rigidity in a functionally critical hinge region impairs the dynamic allosteric residue coupling, leading to a dysfunctional protein [35]. Our data also suggests a link between DCI and the severity of disease mutations. The median %DCI for mutation sites marked as “severe” was 69.6%. In comparison, mutations marked as “mild” had a median of 56.6% (*P* = .045). This further supports the idea that positions exhibiting higher dynamic coupling to the active site have a greater impact on protein function.

**Fig 3.**
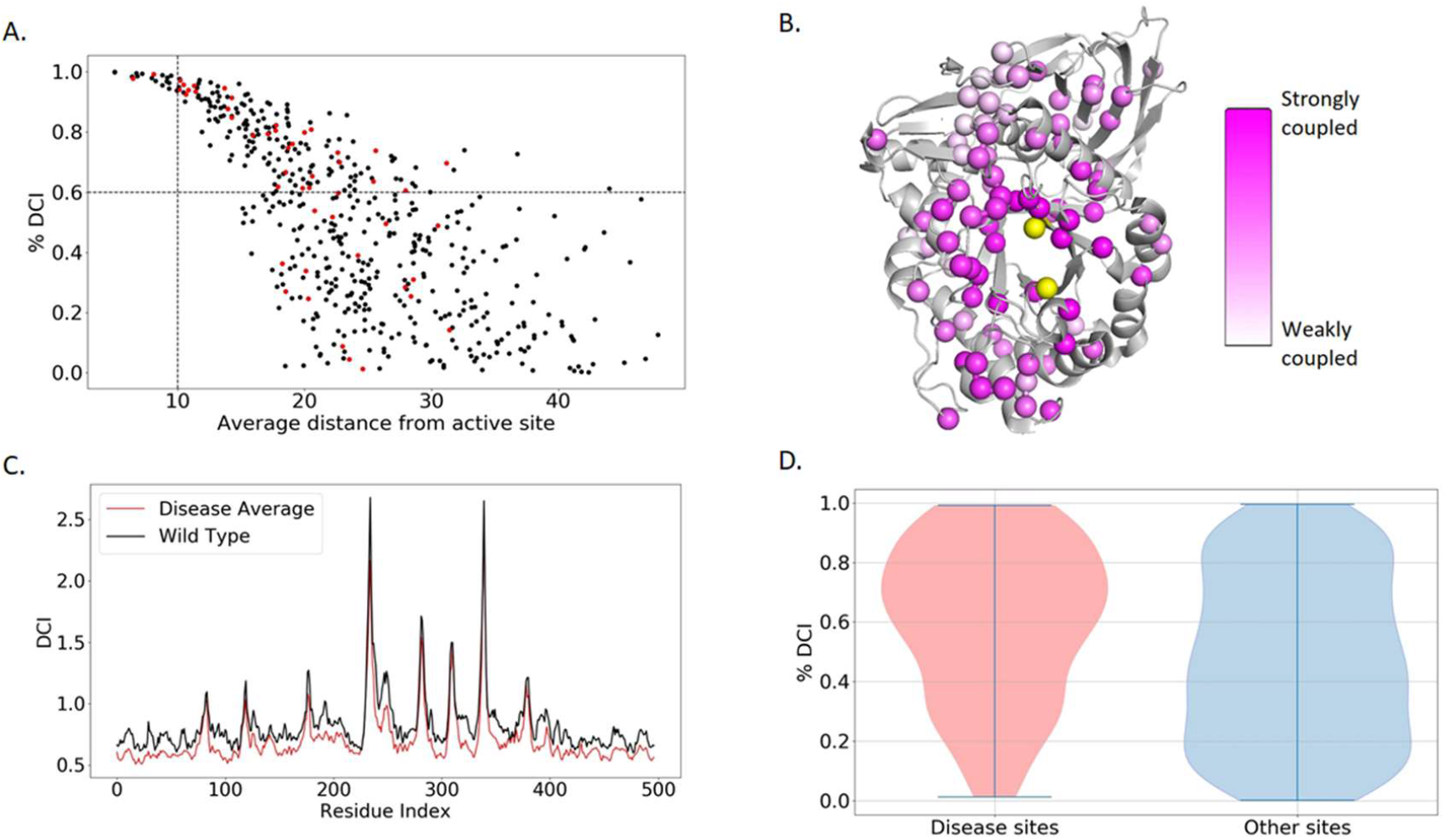
DCI information for residues within GCase. (A) Scatter plot with dividers at %DCI = 60 and distance = 10 Å. The upper right quadrant contains DARC sites which can affect the active site without direct interaction. Red dots indicate severe disease mutations. (B) A ribbon diagram showing known mutation sites of GCase (represented as pink-colored dots) and the degree of coupling to the active site (delineated by the color gradient, where darker and lighter shades correspond to strongly coupled and weakly coupled, respectively). (C) Average disease DCI profile compared to the wild type. In general, we observe a global loss of coupling to the active site (D) Violin plots showing that disease mutations are generally located at sites that have higher DCI with the active site.

### Principal component analysis of DFI aligns with experimentally determined catalytic activity

As explained above, DFI profiles provide information about the dynamic function of residues throughout the protein. At the same time, DARC sites are coupled to the active site despite having no direct contact. By clustering the DFI values of DARC sites for each simulated GCase variant (Fig 4), we observe that the wild type and neutral variants (functional enzymes based on in-vitro assays) are grouped, and many of the tested proteins creating “dead enzymes” (i.e., total loss of function) are grouped as well. Liou et al.[34] used the specific activity of cross-reacting immunological material (CRIM_SA) values to estimate the catalytic rate constants (k_cat_), thereby giving experiment-based estimates on the functionality of these variants. The fact that variants with higher CRIM_SA values are clustered together, as are variants with low CRIM_SA values, suggests a direct correlation between DFI profiles and CRIM_SA and therefore a direct correlation between DFI and protein function.

**Fig 4.**
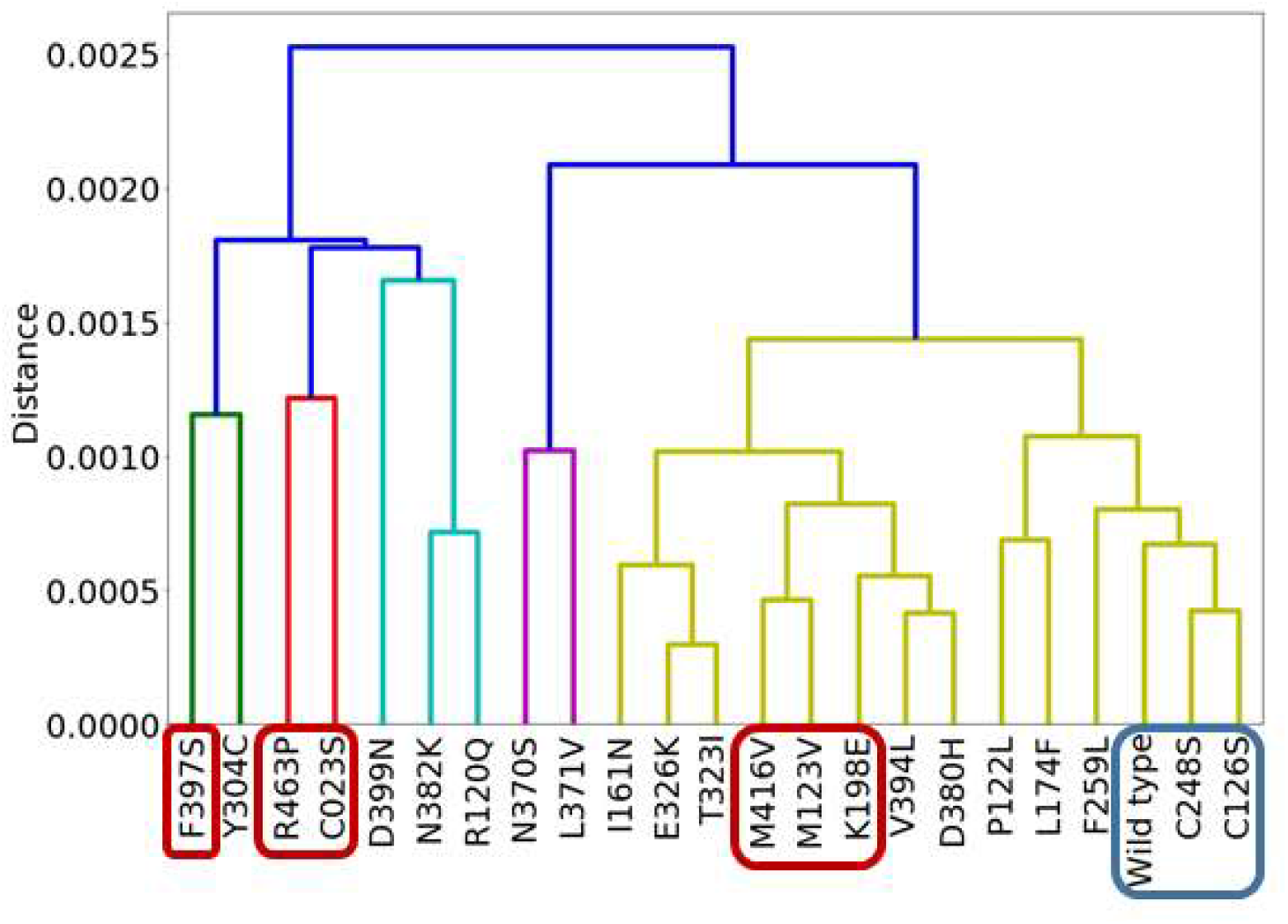
Dendrograph showing clusters of GCase variants based on the DFI of DARC si. The variants for which experimental data is available show dead enzymes and functional enzymes clustered within their own groups. Variants with CRIM_SA values of 0.3 to 1.0 are shown in blue, while variants with CRIM_SA values of 0.06 to 0.1 are shown in red. CRIM_SA data for other variants are unknown. Higher CRIM_SA values suggest superior enzyme function.

### A proteome-wide analysis reveals disease-associated mutations are abundant at DARC sites and hinge sites

After investigating GCase alone, we expanded the study to include 144 human enzymes containing a total of 1024 amino acid variants (433 neutral and 591 disease-associated). The dataset was also used in our previous work [38] incorporating the HumVar data set [39] and sequences with both a high query coverage (>80%) and sequence identity (>80%) selecting only the proteins available in the protein data bank [40]. Additionally, these protein structures had already been modeled, including any missing residues, using the Modeller software package [41].

As best illustrated in Fig 5a, the DCI distribution of the positions with disease-associated variants shows an opposite trend compared to neutral variants. Generally, disease mutations are more likely to occur at positions highly-coupled to the active site. In contrast, neutral mutations are more likely to occur at positions that are less coupled. Of the mutations in this ensemble, 82% occur farther than 10 Å from the active site, necessitating communication through a network of interactions to affect that area of the protein.

**Fig 5.**
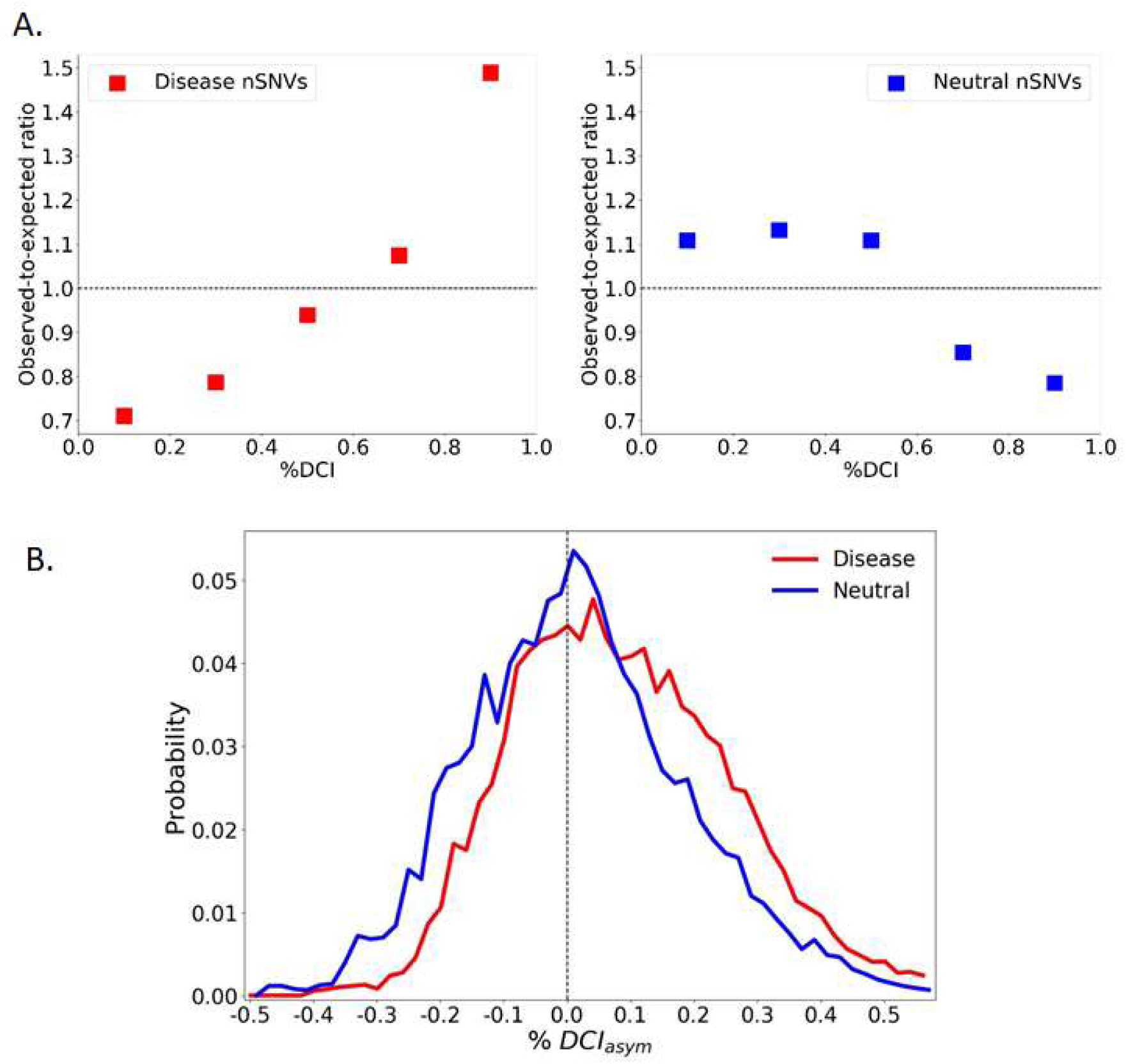
%DCI and asymmetry for 144 protein ensemble. (A) Observed-to-expected ratios reveal that throughout 144 proteins, the number of disease mutations is more than expected at residues that are highly-coupled to the active site, whereas the number of neutral mutations are less than expected at these locations. Above the ratio equal to 1, the mutation type occurs more often than the null expectation, that rates remain the same regardless of DCI. Below the ratio of 1, the mutation does not occur as often as expected. (B) Comparison of %DCI_asym_ of sites associated with neutral and disease variants. The distributions show a contrast where disease variant sites tend to exhibit more positive values (*P*<.001), suggesting that the active site dominates the coupling. Neutral sites on the other hand tend to give more negative asymmetry values, suggesting that the mutation site dominates. A moving average was used to visually smooth the distribution.

DCI specifically quantifies the coupling between individual positions and, as such, DCI values depend explicitly upon the positions selected for analysis. However, these pairwise interactions are not always symmetric. An interaction network may be formed such that residue perturbations may be felt more strongly in one direction than the other. If we find the difference in the DCI values between two residue positions that are not directly interacting (i.e., in spatial contact), we get a better understanding of the dynamic allostery relationship between two residues. This difference, called DCI asymmetry, provides directionality to long-distance coupling, thereby suggesting a causal relationship. In any given protein, every amino acid position has a unique network of direct, local interactions that give rise to a unique network of highly coupled partner positions [9,24,42] and heteronegeity in a 3-D network of interactions. Thus, for a particular pair of coupled amino acids (*i* and *j*), their unique network constraints differentiate the coupling of *i* to *j* from the coupling of *j* to *i*. Thus, we used the wild-type structures of our enzymes to calculate i) %DCI_*i*_, how strongly the position of each mutation is coupled to each active site position, ii) %DCI_*j*_, how strongly each active site position is coupled to the position of each mutation. From these, we calculated iii) “%DCI_asym_” from (%DCI_*i*_–%DCI_*j*_) to assess the asymmetry in coupling.

Among our protein ensemble, we see a slight pattern emerge, where the interaction between disease mutation sites and active sites is generally more dominated by the active sit. In contrast, the interaction between neutral mutation sites and active sites is usually dominated by the mutation site. (Fig 5b). This is indeed in agreement with our earlier findings of LacI variants [9], in which the substitutions on the sites where functional sites dominate the communication most often end up with a function loss.

## Discussion

Allostery was proposed as an important biophysical mechanism for protein function, which has led some to proclaim that allostery constitutes “the second secret of life,” with the genetic code constituting “the first secret of life” [43].

Laboratory-directed evolutionary studies also highlight the emergence of mutations far from the active site [24,44,45]. These distal sites play a critical role in functional evolution, particularly in the emergence of novel functions. Yet, these distal mutation sites challenge enzyme design, as it is difficult to predict them in advance [24,46– 48]. Likewise, resurrected ancestral protein studies also reveal that mutations distal from the active site are necessary for functional evolution. An example is the emergence of red color from a green ancestor in a close relative of Green Fluorescence Protein (GFP). This protein needs a minimum of 12 mutations and one deletion to convert from green to red color with high efficiency. A majority of the mutations are far from the chromophore. While the flexibility of the mutational sites does not change, in allosteric response to these mutations, both rigidification and increased flexibility occur for regions of the fold widely separated in the 3-D structure of the proteins, accommodating required flexibility for red photoconversion. These synergistic effects allow catalysis to proceed as desired and function without mutations of catalytic residue positions while maintaining fold stability and quaternary structure [18].

Here we also observe that disease-associated (i.e., function altering) mutations follow the same pattern. As neutral and disease-associated missense variants provide the best opportunity to explore the molecular principles of how genetic variations shape phenotypic changes, we observed the same principle of dynamic allostery such that functions become altered through distal mutations while conserving the amino-acid sequence of catalytic residues. We have found that the disruption of the allosteric dynamics with functionally-important sites in a protein is a mechanistic explanation for many missense variants associated with diseases and other biological phenotypes. The patterns of dynamic coupling with the active sites are different for disease and neutral phenotypes for missense mutations that occur at spatially-distant positions to functional (active) sites. Specific analysis of GCase proteins also provides evidence of the same mechanism observed in resurrected studies. These distal mutations allosterically modify flexibility profiles of different sites, leading to a change in function.

This finding also suggests that rather than affecting only protein stability, the disruption of ligand binding, or both, the allosteric dynamic coupling and stability explain how a large fraction of disease-associated variants impair protein-ligand function or enzymatic activity [6,7,12]. A high-throughput functional assay of over 2,000 variants also show that only a minority of mutations led to a decrease in protein stability [4]. Thus, our findings align with neutral theory, as mutations on functionally important catalytic sites must have been eliminated as negative selection due to critical functional loss. On the other hand, the distal mutations remotely fine-tune the native state ensemble to modify function without interfering with folding/folding stability.

We are in the era of rapid development of next-generation methods for whole-genome, whole-exome, and targeted sequencing that has produced an unprecedented amount of data. Among all the genetic variation data, the most commonly observed variants are missense, and identifying the missense variants with pathogenic effects that contribute to disease or drug sensitivities is the primary goal of 21^st^-century genomic analysis and phylomedicine. As stated in a review of allostery by Liu and Nussinov [43], uniting the genetic code, which constitutes “the first secret of life,” and allostery, “the second secret of life,” could reveal a generalized disease mechanism and allow for the discovery of novel drugs, as well as blueprints for innovative personalized treatment methods.

## Methods

### Dataset

A total of 144 individual monomeric protein structures from the Protein Data Bank (PDB)[40] were collected from a BLAST search of sequences with requirements of ≥80% sequence identity and ≥80% query coverage to ensure only structures that could be accurately mapped to human variation data were included. Human genetic variations were obtained from the HumVar, and HumDiv databases [37] with 1024 amino acid variants, where 433 were neutral and 591 were deleterious.

### Determining catalytic sites

The catalytic sites were gathered from the Catalytic Site Atlas (CSA) database [49], which identifies the residues directly involved in catalyzing the reactions of enzymes. Since these residues are critical for protein function, they were used as input into our dynamic coupling index (DCI) metric. The entries in the CSA were either “original entries” derived from the literature itself or “homology entries” based on sequence comparison with the literature-based original entries. In either case, the catalytic sites purported by the CSA should accurately represent functional sites on the protein. Our dataset contained 144 enzymatic proteins that mapped to entries in the CSA database.

### Calculating functional-dynamics profiles

Dynamic flexibility index (DFI) quantifies the dynamic stability of a given position. It measures the resilience of a position to perturbations initiated at positions in the protein distal to the residue in question, but to which it is linked via structurally encoded global dynamics. Therefore, DFI profiles provide important information about protein function. Namely, residues that exhibit very low DFI scores (DFIs) do not show large amplitude fluctuations in response to random Brownian kicks but rather transfer the perturbation energies throughout the chain in a cascade fashion; examples of low *DFI* residues are those in hinge regions. Hinges are parts of the protein which are generally rigid. At the same time, they do not exhibit a high fluctuation response to perturbations but transfer these perturbations to the rest of the protein. Like hinges on a door, they stand still, providing an anchor point for other parts to move around.

The method for obtaining the dynamic flexibility index (DFI) is based on the perturbation response scanning (PRS) method [50], in which the C-alpha atom of each residue in the protein is modeled as a node in an elastic network model (ENM). The interaction between each node is modeled by a harmonic potential with a distance-dependent spring constant [50,51]. A small perturbation in the form of an external random force (i.e., Brownian kick) is sequentially applied on each node in the network, and the perturbation response of all nodes is recorded according to linear response theory as

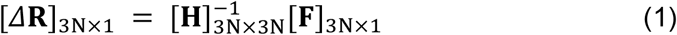

where **F** is the external random force, **H**^-1^ is the inverse Hessian, and Δ**R** is the positional displacement of all *N* nodes in three dimensions.

However, ENM is a coarse-grained model. To improve the accuracy of this model and allow sensitivity to mutations, the hessian inverse can be replaced with the covariance matrices obtained from molecular dynamics simulations.

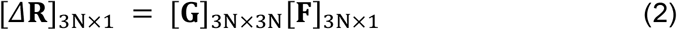

Here, G is the covariance matrix containing the dynamic properties of the system. The covariance matrix contains the data for long-range interactions, solvation effects, and biochemical specificities of all types of interactions.

Each perturbation is performed in ten different directions to ensure an isotropic response. The perturbation is repeated for every node in the network, and the positional displacements Δ**R** of each node are stored in a perturbation matrix **A** given by

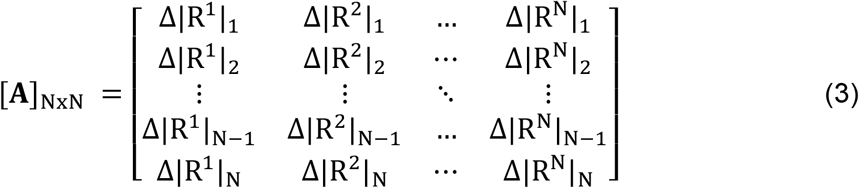

where 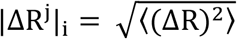 is the magnitude of the positional displacement of each residue *i* in response to a perturbation at residue *j*. The DFI score of residue *i* is defined as the sum of the total displacement of residue *i* induced by a perturbation on all residues, which is computed by taking the sum of the *i-*th row of the perturbation matrix **A**,

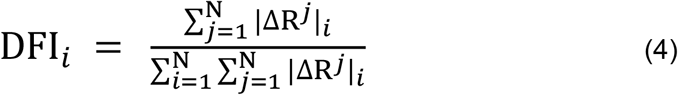

where the denominator is the total displacement of all residues, used as a normalizing factor.

Recently, we have extended this method to identify allosteric links or dynamic coupling between any given residue and functionally important residues by introducing a new metric, the *dynamic coupling index* (DCI) [35]. The DCI metric can identify DARC sites, which are distal to functional sites but control them through dynamic allosteric coupling. This type of allosteric coupling is important; sites with strong dynamic allosteric coupling to functionally critical residues (DARC sites), regardless of separation distance, likely contribute to the function. Thus, a mutation at such a site can disrupt the allosteric dynamic coupling or regulation, leading to functional degradation. As defined, DCI is the ratio of the sum of the mean square fluctuation response of the residue *i* upon functional site perturbations (i.e., catalytic residues) to the response of residue *i* upon perturbations on all residues. DCI enables us to identify DARC site residues, which are more sensitive to perturbations exerted on residues critical for function. This index can be utilized to determine residues involved in allosteric regulation. It is expressed as

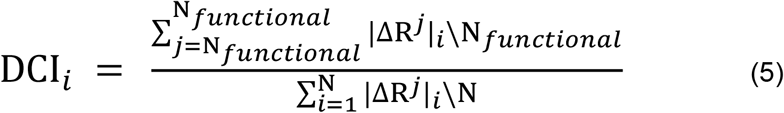

where |ΔR^*j*^|_*i*_ is the response fluctuation profile of residue *i* upon perturbation of residue *j*.

The numerator is the average mean square fluctuation response obtained over the perturbation of the functionally critical residues *N*_functional_. The denominator is the average mean square fluctuation response over all residues. As discussed below, the DFI and DCI profiles can also be computed using the covariance matrix obtained from Molecular Dynamics simulations rather than the inverse Hessian obtained from the elastic network model.

### Molecular Dynamics Simulations

To compute the DFI and DCI profiles of each missense variant, we first performed MD simulations to obtain the native ensemble of each variant and then applied our analysis. Molecular Dynamics (MD) simulations were performed using the AMBER 16 MD package [52]. Simulations were run using the Amber14SB forcefield. The TIP3P [53]water model was used for solvation. The pmemd.cuda [54] executable of the AMBER14 package was used for GPU acceleration. All simulations were run for 1000ns.

To obtain dynamics for each variant, each trajectory was divided into 50ns windows. The covariance matrix **G** for each window was extracted instead of the inverse hessian in the ENM (as in Equation 1), and used to calculate the corresponding DFI and DCI profiles as

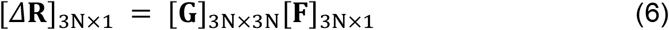

The DFI and DCI profiles calculated from the last half of the trajectory (the final 500ns) were averaged to calculate an average DFI and DCI profile.

We further investigated the change in dynamics upon mutation compared to the wild type structure using ΔDFI and ΔDCI. The delta-DFI (ΔDFI) profile was calculated as

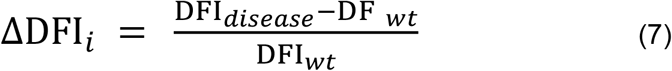

Where DFI_*disease*_ is the dynamics profile for the mutated protein structure and DFI_*wt*_ is the dynamics profile for the wild-type structure. Similarly, the delta-DCI (ΔDCI) profile was calculated as

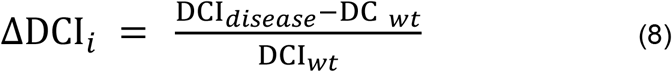

One additional tool we use is DCI asymmetry, which measures preferential information transfer through asymmetric dynamic coupling. Simply put, the coupling asymmetry between positions i and j can be calculated as

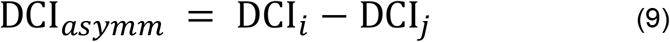

## Supporting information

Enzyme ensemble information

## Supporting Information

**S1 Fig.**
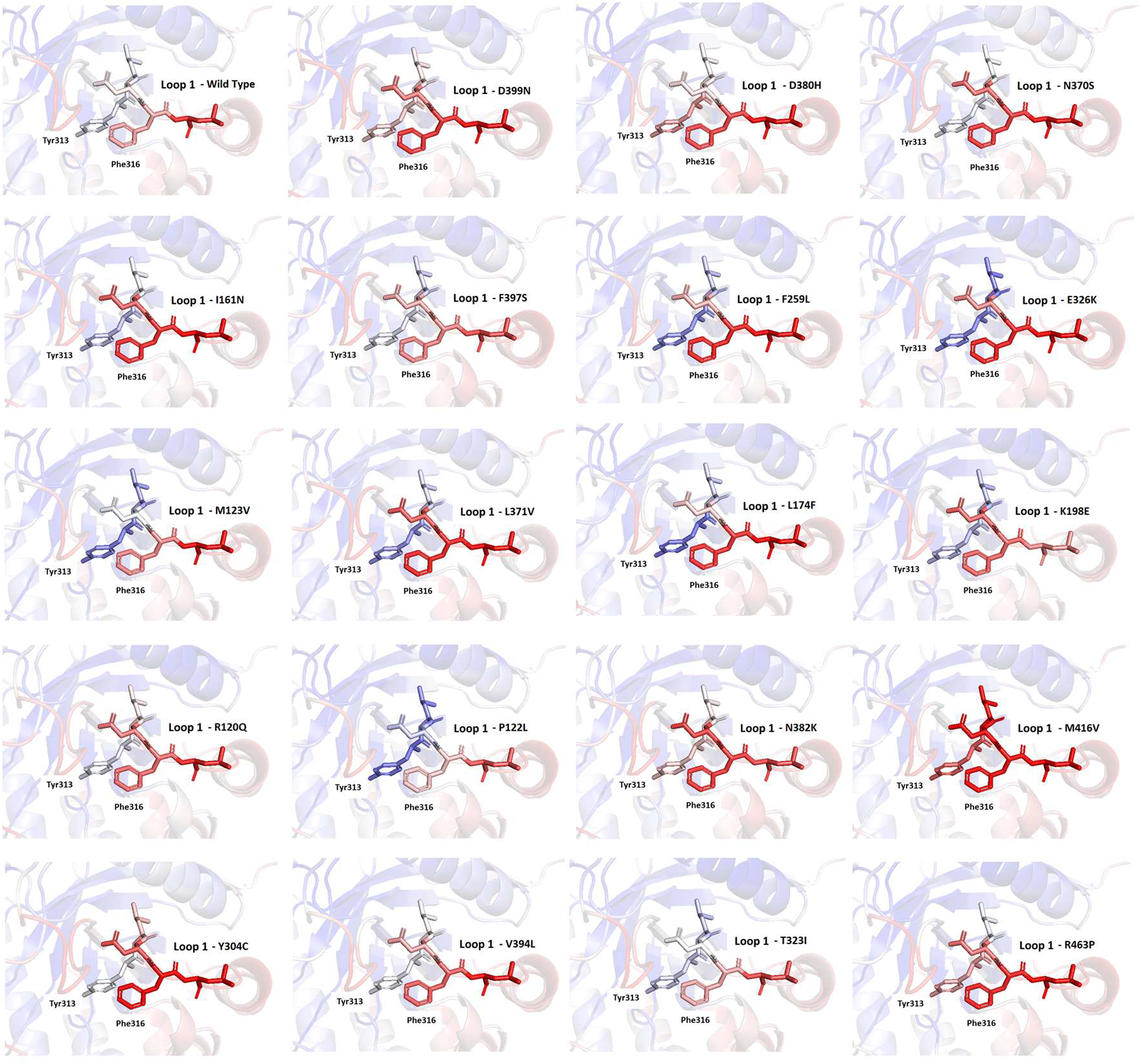
Stick diagrams of loop 1 which are colored corresponding to their DFI for 19 different disease variants. DFI here is a color code within a spectrum of red-white-blue where red shows the highest, and blue shows the lowest flexible sites.

## Notes

https://github.com/SBOZKAN/DFI-DCI

### Competing Interest Statement

The authors have declared no competing interest.

